# Hemoglobin Rothschild: Structural Rationalization of Decreased O_2_ Affinity as a Consequence of a β37(C3)Trp→Arg Mutation

**DOI:** 10.1101/2024.06.06.597804

**Authors:** Mohammed Al-Seragi

**Affiliations:** University of British Columbia

## Abstract

Hemoglobin Rothschild is characterized by a β37(C3)Trp→Arg mutation that severely impairs wildtype hemoglobin function. This mutation has previously been documented to diminish conformational cooperativity, and thereby uppercut oxygen affinity. While the mutation is known to have direct implications on the hinge region at the *α*_1_*β*_2_ interface, the immediate and indirect manifestations of this mutation have not been rendered using high-resolution molecular visualization software. Further unexplored is whether low O_2_ affinity in *Hb*_*R*_ is an outcome of a stabilized, unliganded, tetrameric T-state, a liganded, dimerized R-state deprived of quaternary enhancement, or a combination of both.

Herein, PyMOL is used to rationalize the structural artifacts of the Rothschild variant that govern decreased O_2_ affinity via a stabilized, tetrameric T-state *Hb*_*R*_, and decreased O_2_ affinity via *Hb*_*R*_ dimerization and loss of cooperative binding in the R-state. Molecular docking simulations were then performed to determine on what grounds O_2_ affinity is most attenuated. The result shows that, at the 95% confidence level, reduced O_2_ affinity in *Hb*_*R*_ is just as much an outcome of a stabilized tetrameric T-state as it is a dimerized R-state lacking quaternary, subunit cooperativity. The work described here builds a statistical framework to accommodate further, pair-wise comparison of low O_2_ affinity hemoglobin variants to build intuition on which primary sequence mutations pose the largest clinical consequences.

## Introduction

Wildtype Hemoglobin (*Hb*_*WT*_) is a tetrameric, red blood cell protein composed of two identical alpha (*α*) and beta (*β*) subunits that play a key role in O_2_ transport. Bound to each subunit is a hydrophobic, heme prosthetic group featuring a tetrapyrrole-coordinated Fe^2+^ center that accommodates an O_2_ molecule in a sequestered binding pocket. The wildtype, quaternary structure can interconvert between a low O_2_ affinity, *T-state* conformation, and a high O_2_ affinity, *R-state* conformation: a transition marked by the first O_2_ molecule (*O*_*2*_^*F*^) to bind any of the four subunits, changing orientation of the F-helix (1). The interdependence and allosteric regulation between subunit conformations is known as *cooperative binding* and forms the backbone of O_2_ deposit and uptake between the lungs and all parts of the body.

Gacon et al. (2) were the first to report a hemoglobin variant, coined *Rothschild* (*Hb*_*R*_), characterized by a β37(C3)Trp→Arg mutation, resulting in destabilization of the *α*_1_*β*_2_ contact region modulating the T→R state transition (3), and consequently, dimerization of the protein upon binding *O*_*2*_^*F*^ (1-7). *Hb*_*R*_ has since been characterized by its markedly high *P*_50_ value and low O_2_ uptake (*Figure 1*), owing to non-covalent, T-state-stabilizing interactions in the unliganded tetramer, and an R-state, liganded dimer that no longer benefits from cooperative binding, reminiscent of the Hirose β37(C3)Trp→Ser, Peterborough β111(G13)Val→Phe, and Stanmore β111(G13)Val→Ala variants (2, 8). With arterial O_2_ saturation as low as 81% relative to *Hb*_*WT*_ at 96%, subjects of *Hb*_*R*_ tend to develop clinical cyanosis and anemia (3, 9).

**Figure 1:**
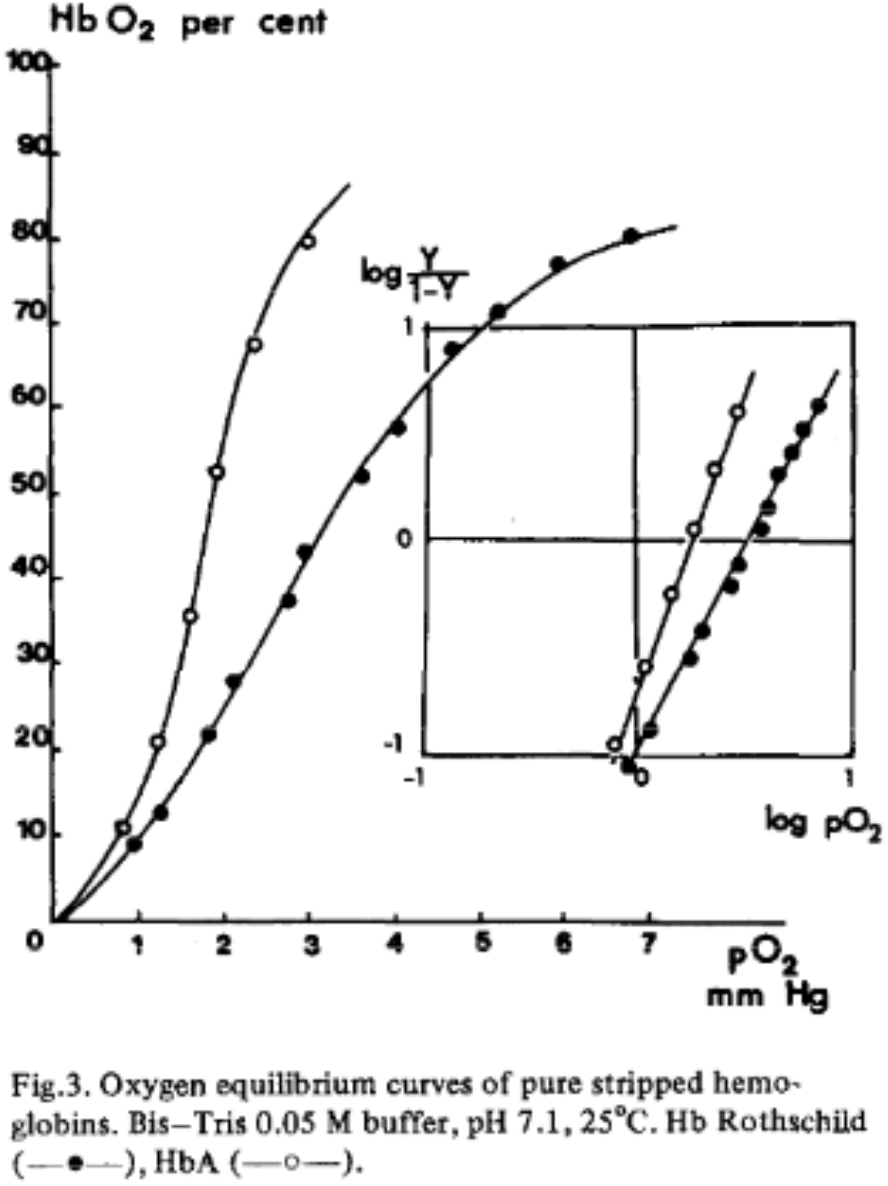
O_2_ binding curve for *Hb*_*R*_, with characteristically low O_2_ saturation across all pO_2_ values, from (2).

While the consequences of *Hb*_*R*_ have been studied extensively, the two modes of decreased O_2_ affinity in this protein have yet to be evaluated side-by-side. As such, this paper aims to qualitatively understand structure-function relationships in *Hb*_*R*_, while quantitatively determining if low O_2_ affinity in *Hb*_*R*_ is an outcome of a stabilized, unliganded, tetrameric T-state, a liganded, dimerized R-state deprived of quaternary enhancement, or a combination of both.

### Decreased O_2_ affinity via a stabilized, tetrameric T-state *Hb*_*R*_

Decreased O_2_ affinity can be attributed to the structural features of unliganded *Hb*_*R*_ adding tetrameric T-state stabilization. A consequence of β37(C3)Trp→Arg is the arrangement of additional water molecules and a Cl^-^ bind site in the absence of the replaced tryptophan ring (5). β37(C3)Arg forms a salt bridge with the Cl^-^ counterion while participating in additional hydrogen bonds not present in *Hb*_*WT*_. *Hb*_*R*_ benefits further, T-state stabilization by involving *α*_1_94(G1)Asp, *α*_1_94(G2)Pro, and *β*_2_102(G4)Asn in polar interactions with the same water molecule oriented at the mutation site, strengthening linkages at the *α*_1_*β*_2_ interface (*Figure 2*). While Bohr-like in effect, the salt-bridge between β37(C3)Arg and Cl^-^ is independent of pH (3), thereby reducing O_2_ affinity in unliganded, tetrameric *Hb*_*R*_ by 10 fold relative to *Hb*_*WT*_ (7).

**Figure 2:**
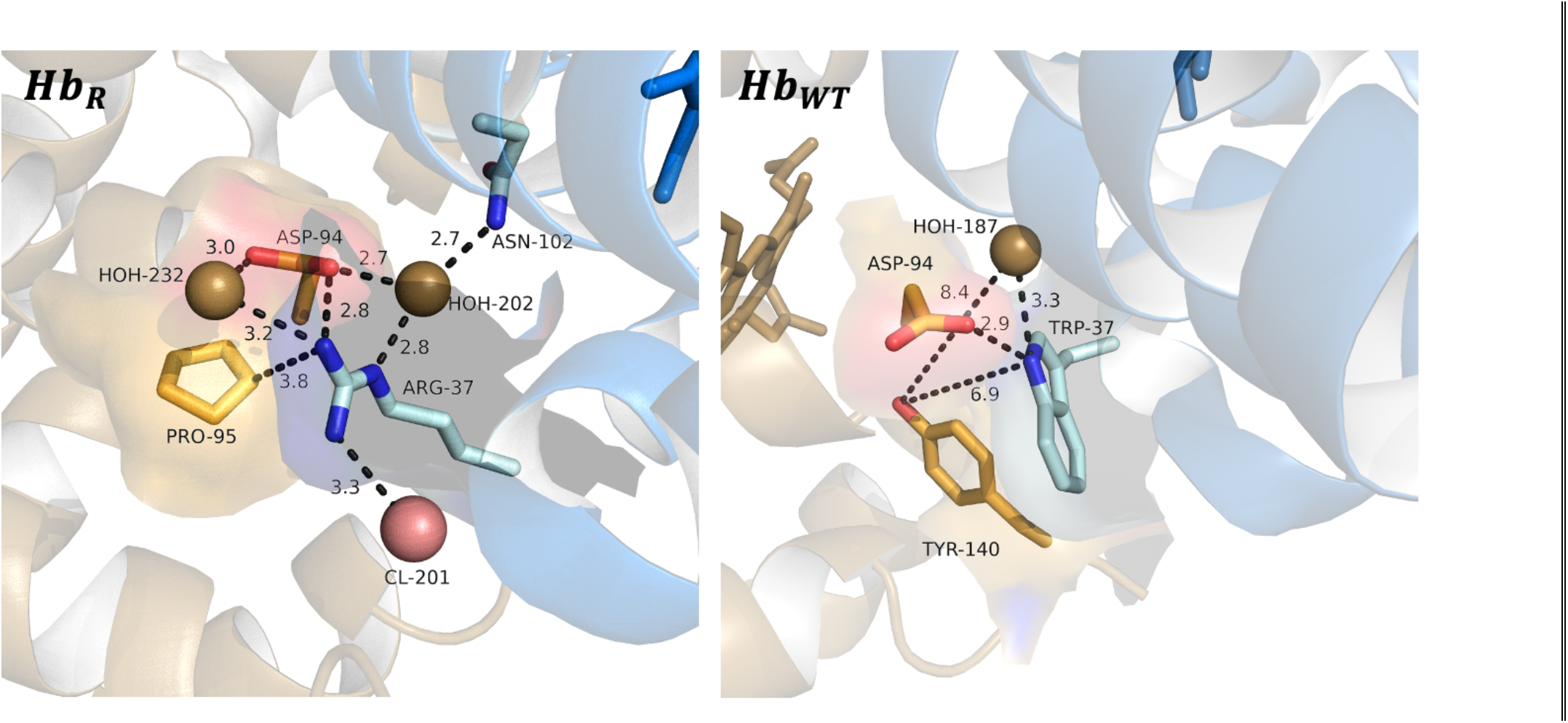
Non-covalent interactions in *Hb*_*R*_ (left) and *Hb*_*WT*_ (right) at the *α*_1_*β*_2_ interface where β37(C3)Trp→Arg occurs. The *α*_1_ subunit is colored gold, and the *β*_2_ subunit is colored blue. Dashed lines mark non-covalent interactions, with distances measured in angstroms (Å) – these conventions are maintained through all figures. <u>Note</u> in *Hb*_*R*_ the presence of a Cl^-^ bind site and additional water molecules recruiting interactions with residues not active in *Hb*_*WT*_. Drawn using Pymol.

An outcome of β37(C3)Trp→Arg extending further away from the site of mutation is the shift in the E-helix relative to the *α*_1_*β*_2_ interface (5) which has implications on distal β63(E7)His stabilization of O_2_ bound to the *β*_2_ heme group. Orientation of the distal β63(E7)His in *Hb*_*R*_ is 0.1Å further away from the heme Fe^2+^ relative to *Hb*_*WT*_ (*Figure 3*). A key parameter in O_2_ binding affinity is positioning of the Fe^2+^, and therefore bound O_2_, relative to distal β63(E7)His, with larger distances signifying weaker hydrogen bonding between the epsilon nitrogen proton of β63(E7)His and the second oxygen atom (9-11); this provides stronghold that the β63(E7)His shift imparted by β37(C3)Trp→Arg provides more T-state character. To this end, reduced O_2_ affinity in unliganded, tetrameric *Hb*_*R*_ is just as much an aftereffect of distant changes stabilizing the T-state as it is the ones directly present at the site of mutation.

**Figure 3:**
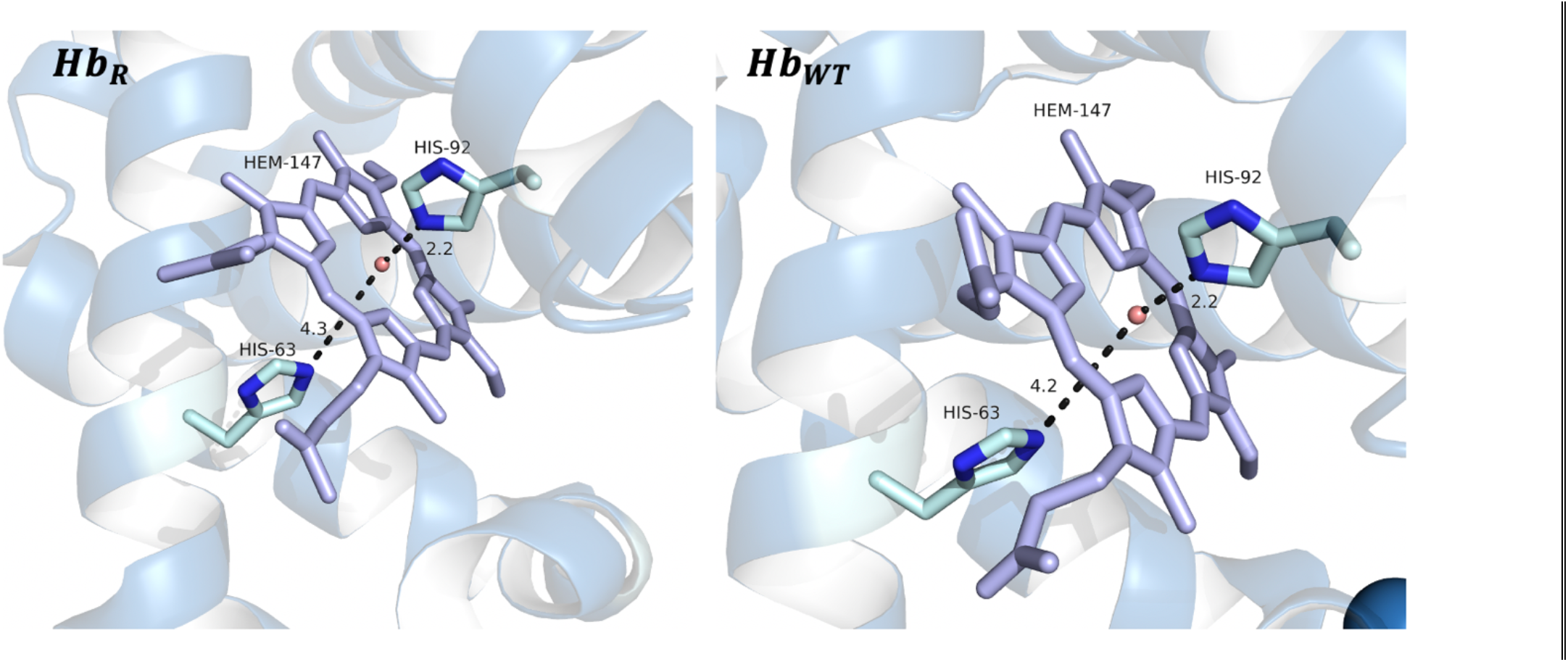
Distances (Å) of distal β63(E7)His and proximal β92(F)His relative to heme Fe^2+^ in *Hb*_*R*_ (left) and *Hb*_*WT*_ (right). <u>Note</u> that the distal His in *Hb*_*R*_ is 0.1Å further away from the *β*_2_ heme. Though not shown in this figure, distal His will hydrogen bond bound O_2_. Drawn using Pymol.

### Decreased O_2_ affinity via *Hb*_*R*_ dimerization and loss of cooperative binding in the R-state

Dimerization of liganded *Hb*_*R*_ tends to be described as the principal cause of reduced O_2_ affinity due to loss of quaternary enhancement in the R-state dimer (4). Of note in tetrameric *Hb*_*R*_ are perturbations in the *hinge region* brought about by bringing the FG corner and the C helix closer together (*Figure 4*). In *Hb*_*WT*_, binding of *O*_*2F*_ triggers the heme ring to flatten, causing the F and G helices to transiently break interactions holding them together, ultimately enabling *α*_1_*β*_2_ and *α*_1_*β*_2_ dimers to rotate and propagate R state configurations across all subunits (1). Additional proximity of the *Hb*_*R*_ F and G helices, however, obstructs this inter-subunit rotation (5), instead driving dimerization and therefore, reduced *Hb*_*R*_ R-state affinity for O_2_. Marked separation of *α*_1_95(*G*2)Pro and *α*_2_96(*G*2)Val in *Hb*_*R*_ was observed as well (*Figure 5*): a shift that has been deemed responsible for the loss of a salt bridge between *α*_1_92(*FG*4)Arg and *β*_2_96(*G*3)Glu at the *α*_1_*β*_2_ contact, again priming the tetramer to dimerize upon binding *O*_*2*_^*F*^ (5). Moreover, loss of space in the *Hb*_*R*_ hinge region, noted in *Figure 4*, may result in sacrifice of the Cl^-^ bind site (5), such that sequestered β37(C3)Arg is left without an anionic binding partner in the hydrophobic, protein interior. Destabilization at this front would also drive dimerization in the pursuit of hydrating the buried, charged residue.

**Figure 4:**
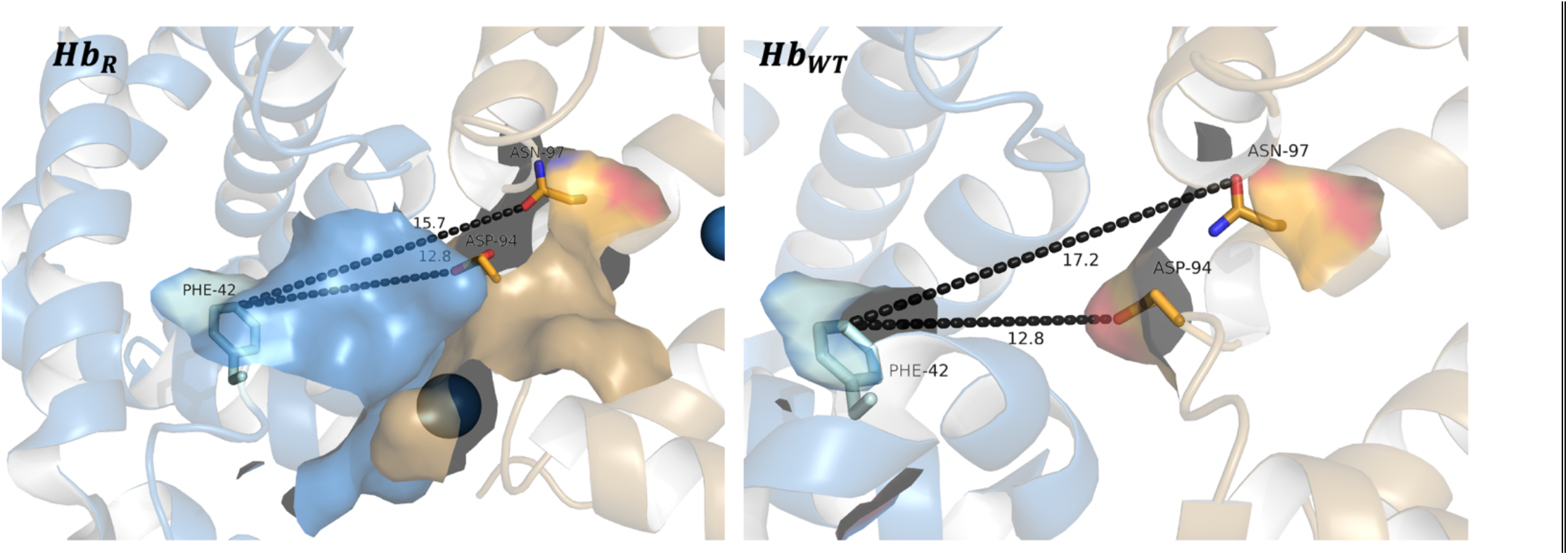
Cleft between *β*_2_ C helix and *α*_1_ FG corner and C helix, termed the *hinge region*, represented as distances (Å) from *β*_2_42(C)Phe to *α*_1_94(*G*1)Asp and *α*_1_97(*G*4)Asn in *Hb*_*R*_ (left) and *Hb*_*WT*_ (right). <u>Note</u> the noticeably smaller hinge region in *Hb*_*R*_ relative to *Hb*_*WT*_. Drawn using Pymol.

**Figure 5:**
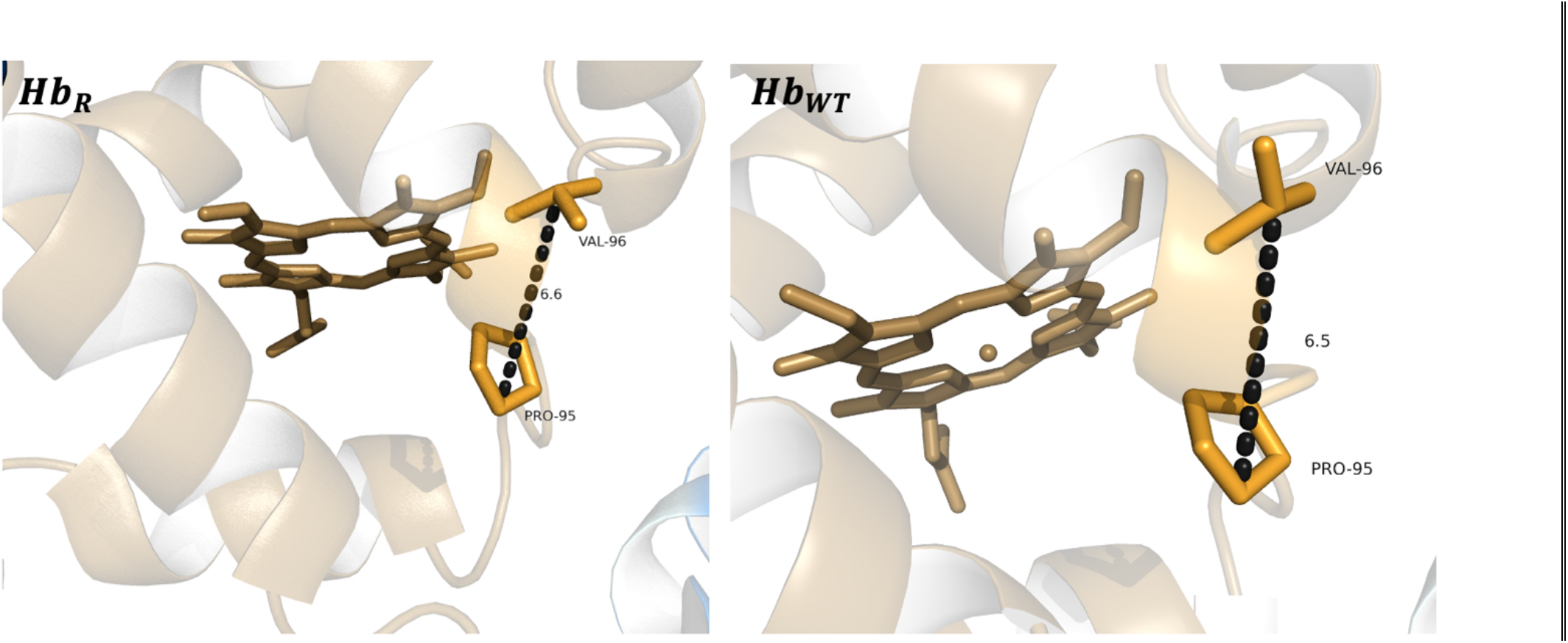
Distance (Å) of non-polar interaction between *α*_1_95(*G*2)Pro and *α*_1_96(*G*2)Val in *Hb*_*R*_ (left) and *Hb*_*WT*_ (right). <u>Note</u> the 0.1Å larger distance in *Hb*_*R*_ relative to *Hb*_*WT*_. Drawn using Pymol.

Other, more ancillary structural cues for *Hb*_*R*_ dimerization and thus, reduced O_2_ affinity, also exist. The apposition of the mutant *β*_2_37(C3)Arg with *α*_1_92(FG4)Arg, for example, causes inter-subunit destabilization at the *α*_1_*β*_2_ contact region via like-charge ionic interactions (*Figure 6*) (4, 5). Furthermore, the lattice of T-state, Cl^-^-bound *Hb*_*R*_, while stabilized by an *β*_2_37(C3)Arg-Cl^-^ salt bridge, is highly incompatible with the conformational transition to the R state (7), and in similar vein to perturbations in the hinge region (*Figure 4*), promotes dimerization upon binding of *O*_*2*_^*F*^. In considering susceptibility of the *Hb*_*R*_ tetramer to dimerize, it is no surprise that “impaired R-state_2_ arguments are the more prevailing rationale for decreased *Hb*_*R*_ O_2_ affinity.

**Figure 6:**
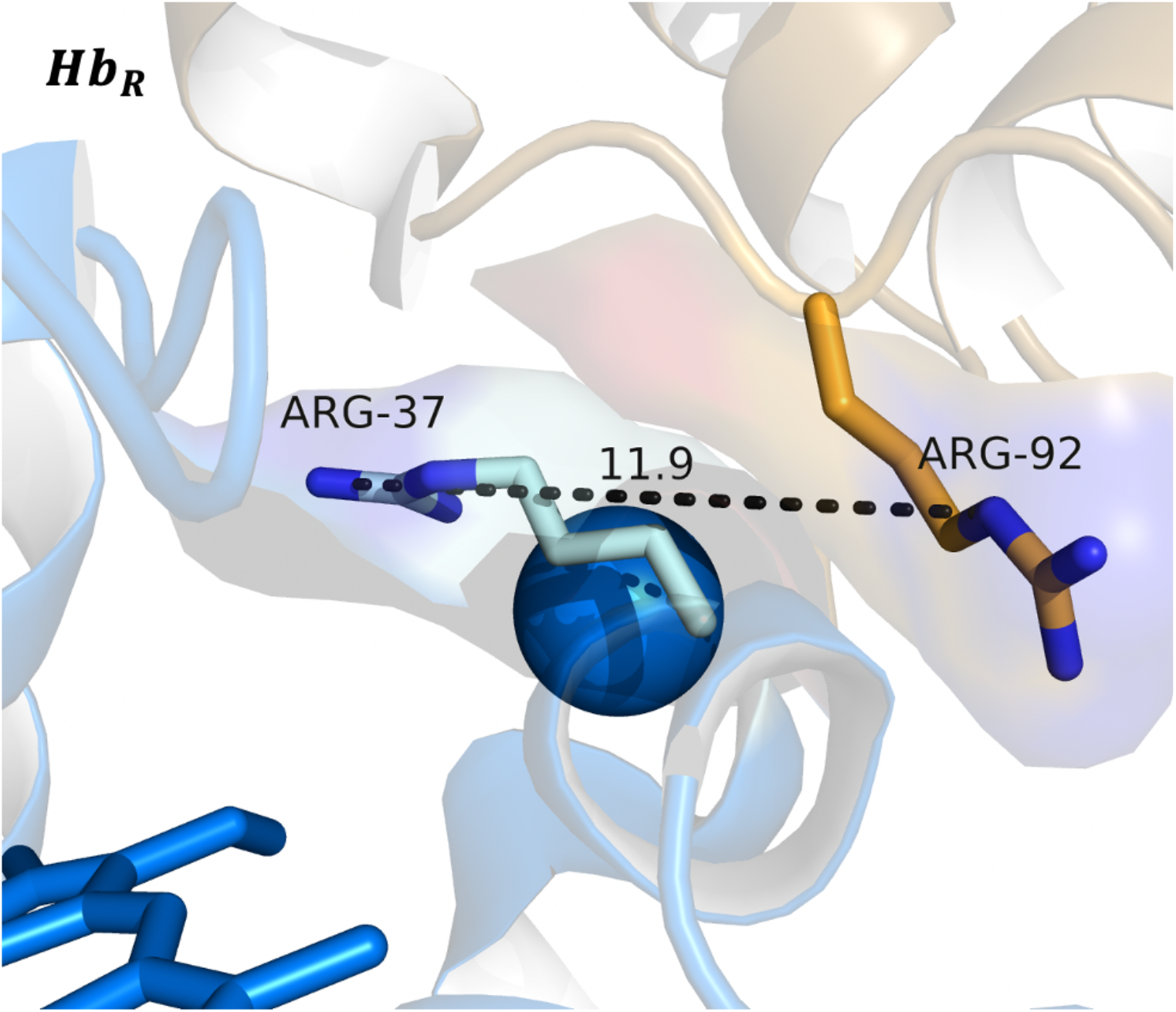
Adjacent, positively-charged *β*_2_37(C3)Arg and *α*_1_92(FG4)Arg contributing instability at the *α*_1_*β*_2_ contact region in *Hb*_*R*_. Drawn using Pymol.

### Decreased O_2_ affinity: Reconciling a stabilized T-state with a dimerized R-state

To date, structural discussion of *Hb*_*R*_, while extensive, has not been used to evaluate the weighted contributions of a stabilized, tetrameric T-state and an impaired, dimerized R state on protein function: *is one mode more significant than the other in reducing Hb*_*R*_ *O*_*2*_ *affinity?*

Investigating this question, UCSF Chimera and SwissDock were used to determine the average biological standard Gibbs free energy of *O*_*2*_^*F*^ binding (Δ*G*^o′^) on the *β*_2_ heme of otherwise unliganded, tetrameric *Hb*_*WT*_ and *Hb*_*R*_ in a computational process known as *docking*. A two-tailed t-test (assuming equal variances) was performed to determine if Δ*G*^o′^ values were statistically different at the *p* = 0.05 level. If Δ*G*^o′^ between both tetrameric *Hb*_*R*_ and *Hb*_*WT*_ are <u>not</u> statistically different, diminished *Hb*_*R*_ affinity for O_2_ is largely an outcome of its dimerization and loss in R-state cooperative binding (**Null Hypothesis**). If Δ*G*^o′^ values <u>are</u> statistically different, the conformational profile in T-state *Hb*_*R*_ pre-dimerization plays a key role in diminished O_2_ affinity by obstruction of *O*_*2*_^*F*^ binding that recruits the T→R state transition across all subunits (**Alternate Hypothesis**). 250 *O*_*2*_^*F*^-*β*_2_ heme binding orientations were produced (*Appendix B*), though only the ones involving the distal β63(E7)His and *β*_2_ heme ring of the O_2_ binding pocket were considered (*Figure 7*).

**Figure 7:**
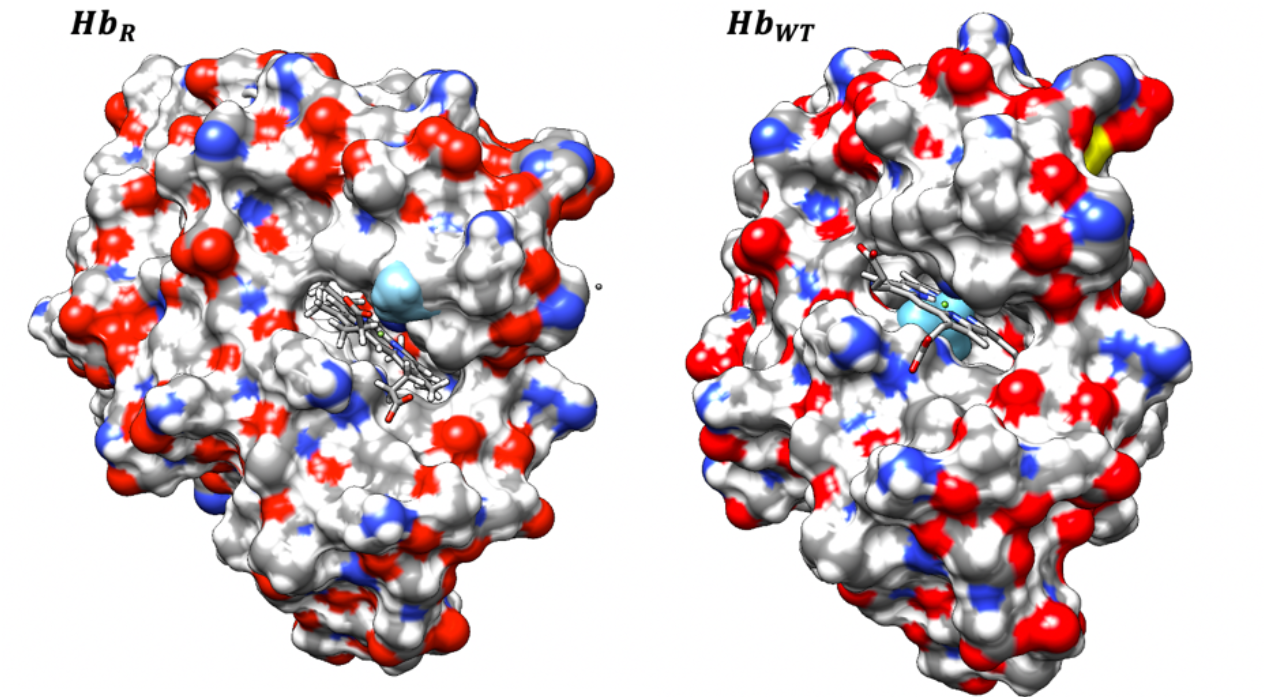
Surface cluster of 30 differently oriented O_2_ molecules (light blue) in the *β*_2_ chain binding pocket of *Hb*_*R*_ (left) and *Hb*_*WT*_ (right). <u>Note</u> that O_2_ molecules aggregate more centrally in the *β*_2_ binding pocket of *Hb*_*WT*_ relative to *Hb*_*R*_ where O_2_ binding is more offset away from the pocket. Drawn using UCSF Chimera.

**Table 1:** Average Δ*G*^o′^ ± 1*σ* of *O*_*2*_^*F*^ Binding (*kj*/*mol*) in the *β*_2_ binding pocket of *Hb*_*R*_ and *Hb*_*WT*_. Details of the t-test are included: degrees of freedom (DOF), *t*_*crit*_, *t*_*exp*_, and conclusion on statistical differences. Calculations can be found in *Appendix A*.

| $\Delta G^{o'} Hb_R - O_2^F$ (kJ/mol) | $\Delta G^{o'} Hb_{WT} - O_2^F$ (kJ/mol) | DOF | $t_{crit}$ | $t_{exp}$ | Conclusion |
| --- | --- | --- | --- | --- | --- |
| $-20.9 \pm 0.4$ | $-21.3 \pm 0.5$ | 29 | 2.1 | 3.4 | $\Delta G^{o'}$ values are statistically different. |

Because *t*_*exp*_ (3.4) > *t*_*crit*_ (2.1), the null hypothesis is rejected, and the alternate hypothesis is accepted, meaning the conformational profile in T-state *Hb*_*R*_ pre-dimerization plays a key role in diminished O_2_ affinity; *i*.*e*. Δ*G*^o′^ *Hb*_*WT*_*-O*_*2*_^*F*^ *is significantly more negative than* Δ*G*^o′^ *Hb*_*R*_*-O*_*2*_^*F*^. This outcome is not aberrant from discussion already made: distal β63(E7)His in *Hb*_*R*_ is 0.1Å further away from the heme Fe^2+^ relative to *Hb*_*WT*_ (*Figure 3*), and this has marked consequences on the thermodynamic stability of bound *O*_*2*_^*F*^ due to weaker hydrogen bonding across larger distances. Also observed in *Figure 7* is the tendency of *O*_*2*_^*F*^ to bind the *β*_2_ heme less centrally, which may be another effect of a perturbed distal β63(E7)His. In any case, this result valuably reinforces that reduced O_2_ affinity in *Hb*_*R*_ is just as much an outcome of a stabilized tetrameric T-state as it is a dimerized R-state lacking quaternary, subunit cooperativity.

## Conclusion

*Hb*_*R*_ is a hemoglobin variant that suffers from reduced O_2_ affinity on the grounds of a β37(C3)Trp→Arg mutation at the *α*_1_*β*_2_ contact region. This can be rationalized through either *stabilized, tetrameric T-state* or *dimerized, impaired R-state* paradigms, but the structural cues in *Hb*_*R*_ lending to either of these low O_2_ affinity states are important to analyze as per statistical analysis performed in this paper. Tetrameric, *Hb*_*R*_ T-state stabilization was discussed to be an outcome of new salt bridges between the mutant β37(C3)Arg and a Cl^-^ counterion, polar interactions from water molecules not present in *Hb*_*WT*_, and repositioning of distal β63(E7)His that detracts *O*_*2*_^*F*^ hydrogen bonding. Conversely, dimerization of *Hb*_*R*_ was largely discussed to be an outcome of arginine apposition and perturbances in the hinge region. While a subject of study for nearly five decades, *Hb*_*R*_ is still the basis of cyanosis and anemia, though it is not the only hemoglobin variant with clinical consequences marked by low hemoglobin O_2_ saturation. Future work can aim to compare *Hb*_*R*_ with variant Hirose (mutated at the same amino acid while also dimerizing) to understand how absence of the β37(C3)Trp residue impacts quaternary structure at-large. Further, pair-wise comparison of low O_2_ affinity hemoglobin variants, such as Peterborough β111(G13)Val→Phe and Stanmore β111(G13)Val→Ala, can also begin to build intuition on which primary sequence mutations pose the largest clinical consequences.

## Appendix A Statistical Test Calculations

An F-test was first performed to determine if the variance between the Δ*G*^o′^ value obtained by *Hb*_*R*_-*O*_*2*_^*F*^ and the *Hb*_*WT*_-*O*_*2*_^*F*^ are statistically different or not.

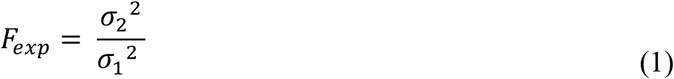

Where *σ*_2_ > *σ*_1_. If *F*_*exp*_ > *F*_*crit*_, then variances are statistically different.

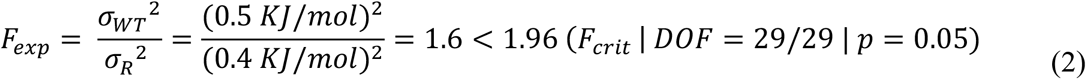

Because *F*_*exp*_ < *F*_*crit*_, the variances between the two measurements are not different at *p* = 0.05. This means a two-tailed t-test tailored for equal variances must be used. This t-test is such that:

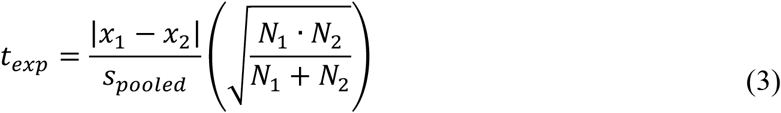

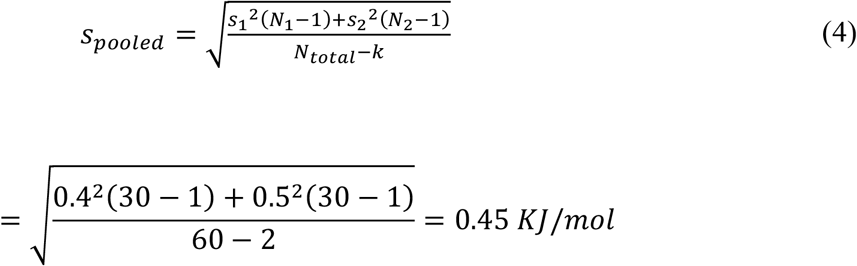

So that,

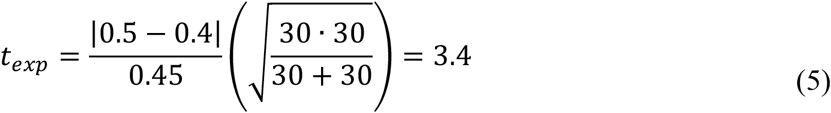

Here, *DOF* = *N* − *k* = 30 − 2 = 28

The results show that *t*_*exp*_ (3.4) > 2.1 (*t*_*crit*_ | *DOF* = 28 | *p* = 0.05). Because *t*_*exp*_ > *t*_*crit*_ at *p* = 0.05, Δ*G*^o′^ *Hb*_*R*_-*O*_*2*_^*F*^ and Δ*G*^o′^ *Hb*_*WT*_-*O*_*2*_^*F*^ are statistically different.

## Appendix B: Raw Data from SwissDock

**Table 2:** Δ*G*^o′^ Values of 250 *O*_*2*_^*F*^-*β*_2_ heme binding orientations, for *Hb*_*R*_. Only values in orange were used to calculate the average found in *Table 1*.

| Cluster | Element | FullFitness (kcal/mol) | Estimated $\Delta G^{o'}$ (kcal/mol) |
| --- | --- | --- | --- |
| 0 | 0 | -855.61 | -5.16 |
| 0 | 1 | -855.61 | -5.16 |
| 0 | 2 | -855.48 | -5.15 |
| 0 | 3 | -855 | -5.11 |
| 0 | 4 | -854.99 | -5.11 |
| 0 | 5 | -853.61 | -4.95 |
| 0 | 6 | -853.61 | -4.95 |
| 0 | 7 | -853.59 | -4.95 |
| 1 | 0 | -854.23 | -5.12 |
| 1 | 1 | -854.02 | -5.01 |
| 1 | 2 | -854.02 | -5.01 |
| 1 | 3 | -854.02 | -5.01 |
| 1 | 4 | -854.02 | -5.01 |
| 1 | 5 | -854.02 | -5.01 |
| 1 | 6 | -854.02 | -5.01 |
| 1 | 7 | -854.02 | -5.01 |
| 2 | 0 | -854 | -4.97 |
| 2 | 1 | -853.86 | -4.96 |
| 2 | 2 | -853.85 | -4.96 |
| 2 | 3 | -853.85 | -4.96 |
| 2 | 4 | -853.65 | -4.91 |
| 2 | 5 | -853.61 | -4.91 |
| 2 | 6 | -853.61 | -4.92 |
| 2 | 7 | -853.46 | -4.89 |
| 3 | 0 | -853.99 | -5.01 |
| 3 | 1 | -853.99 | -5.01 |
| 3 | 2 | -853.32 | -4.89 |
| 3 | 3 | -853.32 | -4.89 |
| 3 | 4 | -853.32 | -4.89 |
| 3 | 5 | -853.32 | -4.89 |
| 3 | 6 | -852.78 | -4.78 |
| 3 | 7 | -852.56 | -4.76 |
| 3 | 8 | -852.52 | -4.86 |
| 3 | 9 | -852.52 | -4.86 |
| 3 | 10 | -852.47 | -4.85 |
| 3 | 11 | -852.07 | -4.76 |
| 3 | 12 | -851.55 | -4.77 |
| 4 | 0 | -853.94 | -5.08 |
| 4 | 1 | -853.94 | -5.08 |
| 4 | 2 | -853.92 | -5.08 |
| 4 | 3 | -853.92 | -5.08 |
| 4 | 4 | -853.92 | -5.08 |
| 4 | 5 | -853.92 | -5.08 |
| 4 | 6 | -853.92 | -5.07 |
| 4 | 7 | -853.92 | -5.07 |
| 5 | 0 | -853.61 | -4.96 |
| 5 | 1 | -853.61 | -4.96 |
| 5 | 2 | -853.6 | -4.96 |
| 5 | 3 | -853.6 | -4.96 |
| 6 | 0 | -853.54 | -5.04 |
| 6 | 1 | -853.53 | -5.04 |
| 6 | 2 | -853.53 | -5.04 |
| 6 | 3 | -853.09 | -4.73 |
| 6 | 4 | -853.09 | -4.73 |
| 6 | 5 | -853.09 | -4.73 |
| 6 | 6 | -852.79 | -4.68 |
| 6 | 7 | -852.7 | -4.67 |
| 6 | 8 | -852.54 | -4.88 |
| 6 | 9 | -852.54 | -4.88 |
| 6 | 10 | -852.54 | -4.88 |
| 6 | 11 | -852.53 | -4.87 |
| 6 | 12 | -852.39 | -4.87 |
| 6 | 13 | -852.39 | -4.87 |
| 6 | 14 | -852.39 | -4.87 |
| 6 | 15 | -852.39 | -4.87 |
| 7 | 0 | -853.51 | -5.14 |
| 7 | 1 | -853.51 | -5.14 |
| 7 | 2 | -853.51 | -5.14 |
| 7 | 3 | -853.5 | -5.13 |
| 8 | 0 | -853.33 | -4.9 |
| 8 | 1 | -853.33 | -4.9 |
| 8 | 2 | -853.33 | -4.9 |
| 8 | 3 | -853.33 | -4.9 |
| 8 | 4 | -853.33 | -4.9 |
| 8 | 5 | -853.33 | -4.9 |
| 8 | 6 | -853.27 | -4.9 |
| 8 | 7 | -853.27 | -4.9 |
| 9 | 0 | -853.32 | -4.79 |
| 9 | 1 | -853.32 | -4.79 |
| 9 | 2 | -853.32 | -4.79 |
| 9 | 3 | -853.32 | -4.79 |
| 9 | 4 | -853.32 | -4.79 |
| 9 | 5 | -853.32 | -4.79 |
| 9 | 6 | -853.14 | -4.78 |
| 9 | 7 | -853.14 | -4.78 |
| 10 | 0 | -853.3 | -4.85 |
| 10 | 1 | -853.3 | -4.85 |
| 10 | 2 | -853.3 | -4.85 |
| 10 | 3 | -853.29 | -4.86 |
| 10 | 4 | -853.29 | -4.86 |
| 10 | 5 | -853.29 | -4.86 |
| 10 | 6 | -853.29 | -4.86 |
| 10 | 7 | -853.28 | -4.86 |
| 11 | 0 | -853.2 | -4.89 |
| 11 | 1 | -853.2 | -4.89 |
| 11 | 2 | -853.2 | -4.89 |
| 11 | 3 | -853.2 | -4.89 |
| 11 | 4 | -853.2 | -4.89 |
| 11 | 5 | -853.2 | -4.89 |
| 11 | 6 | -853.2 | -4.89 |
| 11 | 7 | -853.2 | -4.89 |
| 12 | 0 | -853.2 | -5.25 |
| 12 | 1 | -853.2 | -5.25 |
| 12 | 2 | -853.2 | -5.25 |
| 12 | 3 | -853.17 | -5.25 |
| 12 | 4 | -853.17 | -5.25 |
| 12 | 5 | -853.16 | -5.24 |
| 12 | 6 | -853.07 | -5.23 |
| 12 | 7 | -853.07 | -5.23 |
| 13 | 0 | -853.01 | -4.92 |
| 13 | 1 | -853.01 | -4.92 |
| 13 | 2 | -853.01 | -4.92 |
| 13 | 3 | -853.01 | -4.92 |
| 13 | 4 | -853.01 | -4.92 |
| 13 | 5 | -853.01 | -4.92 |
| 13 | 6 | -853.01 | -4.92 |
| 13 | 7 | -853.01 | -4.92 |
| 14 | 0 | -852.99 | -4.91 |
| 14 | 1 | -852.97 | -4.84 |
| 14 | 2 | -852.97 | -4.91 |
| 14 | 3 | -852.97 | -4.91 |
| 14 | 4 | -852.97 | -4.84 |
| 14 | 5 | -852.97 | -4.84 |
| 14 | 6 | -852.97 | -4.84 |
| 14 | 7 | -852.97 | -4.91 |
| 15 | 0 | -852.88 | -4.85 |
| 15 | 1 | -852.88 | -4.85 |
| 15 | 2 | -852.76 | -4.86 |
| 15 | 3 | -852.76 | -4.86 |
| 15 | 4 | -852.76 | -4.86 |
| 15 | 5 | -852.64 | -4.85 |
| 15 | 6 | -852.64 | -4.85 |
| 15 | 7 | -852.64 | -4.85 |
| 16 | 0 | -852.76 | -4.86 |
| 16 | 1 | -852.76 | -4.86 |
| 16 | 2 | -852.76 | -4.86 |
| 16 | 3 | -852.75 | -4.86 |
| 16 | 4 | -852.75 | -4.86 |
| 16 | 5 | -852.68 | -4.86 |
| 16 | 6 | -852.68 | -4.86 |
| 16 | 7 | -852.68 | -4.86 |
| 17 | 0 | -852.71 | -4.73 |
| 17 | 1 | -852.67 | -4.72 |
| 17 | 2 | -852.67 | -4.72 |
| 17 | 3 | -852.67 | -4.72 |
| 18 | 0 | -852.38 | -4.94 |
| 18 | 1 | -852.36 | -4.93 |
| 18 | 2 | -852.25 | -4.98 |
| 18 | 3 | -851.9 | -4.97 |
| 18 | 4 | -851.89 | -4.97 |
| 18 | 5 | -851.86 | -4.96 |
| 18 | 6 | -851.86 | -4.82 |
| 18 | 7 | -851.05 | -4.91 |
| 19 | 0 | -852.3 | -4.97 |
| 19 | 1 | -852.3 | -4.97 |
| 19 | 2 | -852.3 | -4.97 |
| 19 | 3 | -852.3 | -4.97 |
| 19 | 4 | -852.3 | -4.97 |
| 19 | 5 | -852.3 | -4.97 |
| 19 | 6 | -852.3 | -4.97 |
| 19 | 7 | -852.3 | -4.97 |
| 19 | 8 | -852.12 | -4.94 |
| 19 | 9 | -852.12 | -4.94 |
| 19 | 10 | -852.12 | -4.94 |
| 19 | 11 | -852.12 | -4.94 |
| 19 | 12 | -852.12 | -4.94 |
| 19 | 13 | -852.12 | -4.94 |
| 19 | 14 | -852.12 | -4.94 |
| 19 | 15 | -852.12 | -4.94 |
| 20 | 0 | -852.27 | -5.14 |
| 20 | 1 | -852.21 | -5.14 |
| 20 | 2 | -851.97 | -5.14 |
| 20 | 3 | -851.89 | -5.11 |
| 20 | 4 | -851.89 | -5.12 |
| 20 | 5 | -851.7 | -5.08 |
| 20 | 6 | -851.48 | -5.06 |
| 20 | 7 | -851.45 | -5.05 |
| 20 | 8 | -851.44 | -5.05 |
| 20 | 9 | -851.41 | -5.05 |
| 20 | 10 | -851.12 | -5.02 |
| 21 | 0 | -852.11 | -4.74 |
| 21 | 1 | -852.08 | -4.74 |
| 21 | 2 | -852.08 | -4.74 |
| 21 | 3 | -852.08 | -4.74 |
| 21 | 4 | -852.08 | -4.74 |
| 21 | 5 | -852.08 | -4.74 |
| 21 | 6 | -852.08 | -4.74 |
| 21 | 7 | -852.08 | -4.74 |
| 22 | 0 | -852.1 | -4.76 |
| 22 | 1 | -852.1 | -4.76 |
| 22 | 2 | -852.1 | -4.76 |
| 22 | 3 | -852.09 | -4.76 |
| 22 | 4 | -852.09 | -4.76 |
| 22 | 5 | -852.08 | -4.76 |
| 22 | 6 | -852.08 | -4.76 |
| 22 | 7 | -852.08 | -4.76 |
| 23 | 0 | -852.1 | -4.73 |
| 23 | 1 | -852.1 | -4.73 |
| 23 | 2 | -852.09 | -4.73 |
| 23 | 3 | -851.99 | -4.87 |
| 24 | 0 | -852 | -4.77 |
| 24 | 1 | -852 | -4.77 |
| 24 | 2 | -852 | -4.77 |
| 24 | 3 | -852 | -4.77 |
| 24 | 4 | -851.82 | -4.73 |
| 24 | 5 | -851.82 | -4.73 |
| 24 | 6 | -851.82 | -4.73 |
| 24 | 7 | -851.82 | -4.73 |
| 25 | 0 | -851.93 | -4.91 |
| 25 | 1 | -851.84 | -4.91 |
| 25 | 2 | -851.84 | -4.96 |
| 25 | 3 | -851.83 | -4.96 |
| 25 | 4 | -851.82 | -4.91 |
| 25 | 5 | -851.7 | -4.85 |
| 25 | 6 | -851.62 | -4.85 |
| 25 | 7 | -851 | -4.87 |
| 26 | 0 | -851.92 | -4.77 |
| 26 | 1 | -851.92 | -4.77 |
| 26 | 2 | -851.92 | -4.77 |
| 26 | 3 | -851.92 | -4.77 |
| 26 | 4 | -851.92 | -4.77 |
| 26 | 5 | -851.92 | -4.77 |
| 26 | 6 | -851.92 | -4.77 |
| 26 | 7 | -851.92 | -4.77 |
| 27 | 0 | -851.86 | -4.99 |
| 27 | 1 | -851.86 | -4.99 |
| 27 | 2 | -851.85 | -4.98 |
| 27 | 3 | -851.82 | -4.98 |
| 27 | 4 | -851.7 | -4.97 |
| 27 | 5 | -851.57 | -4.98 |
| 27 | 6 | -851.57 | -4.98 |
| 28 | 0 | -851.76 | -4.75 |
| 28 | 1 | -851.76 | -4.75 |
| 28 | 2 | -851.76 | -4.75 |
| 28 | 3 | -851.76 | -4.75 |
| 28 | 4 | -851.76 | -4.75 |
| 28 | 5 | -851.76 | -4.75 |
| 28 | 6 | -851.76 | -4.75 |
| 28 | 7 | -851.76 | -4.75 |
| 29 | 0 | -851.55 | -4.83 |
| 29 | 1 | -851.55 | -4.88 |
| 29 | 2 | -851.46 | -4.83 |
| 29 | 3 | -851.41 | -4.83 |
| 29 | 4 | -851.41 | -4.87 |
| 29 | 5 | -851.24 | -4.86 |
| 29 | 6 | -850.61 | -4.83 |
| 29 | 7 | -850.54 | -4.83 |
| 30 | 0 | -850.49 | -4.82 |
| 31 | 0 | -850.44 | -4.85 |
| 31 | 1 | -850.42 | -4.85 |
| 31 | 2 | -850.37 | -4.86 |
| 31 | 3 | -850.36 | -4.86 |
| 31 | 4 | -850.2 | -4.83 |
| 31 | 4 | -850.2 | -4.83 |

**Table 3:** Δ*G*^o′^ Values of 250 *O*_*2*_^*F*^-*β*_2_ heme binding orientations, for *Hb*_*WT*_. Only values in orange were used to calculate the average found in *Table 1*.

| Cluster | Element | FullFitness (kcal/mol) | Estimated $\Delta G^{o'}$(kcal/mol) |
| --- | --- | --- | --- |
| 0 | 0 | -806.92 | -5.36 |
| 0 | 1 | -806.84 | -5.36 |
| 0 | 2 | -806.47 | -5.31 |
| 0 | 3 | -806.46 | -5.31 |
| 0 | 4 | -806.17 | -5.28 |
| 0 | 5 | -804.21 | -5.06 |
| 0 | 6 | -804.17 | -5.08 |
| 0 | 7 | -804.17 | -5.08 |
| 0 | 8 | -802.62 | -4.95 |
| 1 | 0 | -806.64 | -5.05 |
| 1 | 1 | -806.63 | -5.02 |
| 1 | 2 | -806.59 | -5.01 |
| 1 | 3 | -806.59 | -5.01 |
| 1 | 4 | -806.34 | -5.03 |
| 1 | 5 | -806.23 | -5.02 |
| 1 | 6 | -806.19 | -5.03 |
| 1 | 7 | -806.19 | -5.03 |
| 2 | 0 | -805.62 | -5.13 |
| 2 | 1 | -805.61 | -5.13 |
| 2 | 2 | -805.59 | -5.13 |
| 2 | 3 | -805.39 | -5.1 |
| 2 | 4 | -805.39 | -5.1 |
| 2 | 5 | -804.89 | -4.96 |
| 2 | 6 | -804.89 | -4.96 |
| 2 | 7 | -804.76 | -4.95 |
| 3 | 0 | -805.15 | -5.02 |
| 3 | 1 | -805.15 | -5.02 |
| 3 | 2 | -805.15 | -5.02 |
| 3 | 3 | -805.14 | -5.02 |
| 3 | 4 | -805.14 | -5.02 |
| 3 | 5 | -805.14 | -5.02 |
| 3 | 6 | -805.03 | -4.99 |
| 3 | 7 | -805.03 | -4.99 |
| 4 | 0 | -805 | -5.04 |
| 4 | 1 | -804.92 | -5.07 |
| 4 | 2 | -804.91 | -5.06 |
| 4 | 3 | -804.86 | -4.99 |
| 4 | 4 | -804.85 | -5.04 |
| 4 | 5 | -804.84 | -4.93 |
| 4 | 6 | -804.82 | -5.04 |
| 4 | 7 | -804.77 | -4.92 |
| 4 | 8 | -804.75 | -5.03 |
| 4 | 9 | -804.72 | -5.03 |
| 4 | 10 | -804.69 | -4.92 |
| 4 | 11 | -804.63 | -4.93 |
| 4 | 12 | -804.63 | -4.93 |
| 4 | 13 | -804.63 | -4.93 |
| 4 | 14 | -804.63 | -4.93 |
| 4 | 15 | -804.63 | -4.93 |
| 4 | 16 | -804.32 | -4.91 |
| 4 | 17 | -804.19 | -4.82 |
| 4 | 18 | -803.93 | -4.78 |
| 4 | 19 | -803.59 | -4.79 |
| 5 | 0 | -804.99 | -5.14 |
| 5 | 1 | -804.99 | -5.14 |
| 5 | 2 | -804.99 | -5.14 |
| 5 | 3 | -803.73 | -5.04 |
| 5 | 4 | -803.73 | -5.04 |
| 5 | 5 | -803.72 | -5.04 |
| 5 | 6 | -803.72 | -5.04 |
| 5 | 7 | -803.72 | -5.04 |
| 6 | 0 | -804.96 | -4.86 |
| 6 | 1 | -804.96 | -4.86 |
| 6 | 2 | -804.96 | -4.86 |
| 6 | 3 | -804.96 | -4.86 |
| 6 | 4 | -804.88 | -4.86 |
| 6 | 5 | -804.88 | -4.86 |
| 6 | 6 | -804.88 | -4.86 |
| 6 | 7 | -804.87 | -4.86 |
| 7 | 0 | -804.94 | -4.96 |
| 7 | 1 | -804.94 | -4.96 |
| 7 | 2 | -804.64 | -4.98 |
| 7 | 3 | -803.87 | -5.03 |
| 7 | 4 | -803.87 | -5.03 |
| 7 | 5 | -803.75 | -4.91 |
| 8 | 0 | -804.94 | -4.96 |
| 8 | 1 | -804.92 | -4.96 |
| 8 | 2 | -804.92 | -4.96 |
| 8 | 3 | -804.92 | -4.96 |
| 8 | 4 | -804.91 | -4.96 |
| 8 | 5 | -804.91 | -4.96 |
| 8 | 6 | -804.78 | -4.94 |
| 8 | 7 | -804.6 | -4.92 |
| 9 | 0 | -804.53 | -5.03 |
| 9 | 1 | -804.53 | -5.03 |
| 9 | 2 | -804.53 | -5.03 |
| 9 | 3 | -804.53 | -5.03 |
| 9 | 4 | -804.53 | -5.03 |
| 9 | 5 | -804.53 | -5.03 |
| 9 | 6 | -804.28 | -5.07 |
| 9 | 7 | -804.28 | -5.07 |
| 10 | 0 | -804.53 | -5.22 |
| 10 | 1 | -804.51 | -5.22 |
| 10 | 2 | -804.51 | -5.22 |
| 10 | 3 | -804.51 | -5.22 |
| 10 | 4 | -804.51 | -5.22 |
| 10 | 5 | -804.51 | -5.22 |
| 10 | 6 | -804.51 | -5.22 |
| 10 | 7 | -804.51 | -5.22 |
| 10 | 8 | -804.51 | -5.22 |
| 10 | 9 | -804.5 | -5.22 |
| 10 | 10 | -803.93 | -5.09 |
| 10 | 11 | -803.93 | -5.08 |
| 10 | 12 | -803.92 | -5.09 |
| 10 | 13 | -803.62 | -4.67 |
| 10 | 14 | -803.62 | -4.67 |
| 10 | 15 | -803.62 | -4.67 |
| 10 | 16 | -802.9 | -4.9 |
| 10 | 17 | -802.86 | -4.9 |
| 10 | 18 | -802.19 | -4.91 |
| 11 | 0 | -804.51 | -5.36 |
| 11 | 1 | -804.51 | -5.36 |
| 11 | 2 | -804.5 | -5.36 |
| 11 | 3 | -804.5 | -5.36 |
| 11 | 4 | -804.5 | -5.36 |
| 11 | 5 | -804.5 | -5.36 |
| 11 | 6 | -804.49 | -5.36 |
| 11 | 7 | -804.49 | -5.36 |
| 12 | 0 | -804.39 | -4.97 |
| 12 | 1 | -804.39 | -4.97 |
| 12 | 2 | -804.39 | -4.97 |
| 12 | 3 | -804.39 | -4.96 |
| 12 | 4 | -804.39 | -4.96 |
| 12 | 5 | -804.39 | -4.97 |
| 12 | 6 | -804.39 | -4.97 |
| 12 | 7 | -804.39 | -4.97 |
| 13 | 0 | -804.35 | -4.9 |
| 13 | 1 | -804.34 | -4.98 |
| 13 | 2 | -804.33 | -4.98 |
| 13 | 3 | -804.33 | -4.98 |
| 13 | 4 | -804.27 | -4.94 |
| 13 | 5 | -804.27 | -4.94 |
| 13 | 6 | -804.27 | -4.94 |
| 13 | 7 | -804.27 | -4.94 |
| 14 | 0 | -804.33 | -4.87 |
| 15 | 0 | -804.21 | -4.98 |
| 15 | 1 | -804.21 | -4.98 |
| 15 | 2 | -804.21 | -4.98 |
| 15 | 3 | -804.21 | -4.98 |
| 15 | 4 | -804.21 | -4.98 |
| 15 | 5 | -804.21 | -4.98 |
| 15 | 6 | -804.21 | -4.98 |
| 15 | 7 | -804.21 | -4.98 |
| 16 | 0 | -804.08 | -4.88 |
| 16 | 1 | -803.79 | -4.86 |
| 16 | 2 | -803.42 | -4.82 |
| 16 | 3 | -803.06 | -4.81 |
| 16 | 4 | -801.77 | -4.61 |
| 16 | 5 | -801.62 | -4.67 |
| 17 | 0 | -803.95 | -4.79 |
| 17 | 1 | -803.95 | -4.79 |
| 17 | 2 | -803.95 | -4.79 |
| 17 | 3 | -803.95 | -4.79 |
| 17 | 4 | -803.95 | -4.79 |
| 17 | 5 | -803.95 | -4.79 |
| 17 | 6 | -803.95 | -4.79 |
| 17 | 7 | -803.95 | -4.79 |
| 18 | 0 | -803.89 | -4.88 |
| 18 | 1 | -803.89 | -4.88 |
| 18 | 2 | -803.89 | -4.88 |
| 18 | 3 | -803.89 | -4.88 |
| 18 | 4 | -803.89 | -4.88 |
| 18 | 5 | -803.89 | -4.88 |
| 18 | 6 | -803.79 | -4.87 |
| 18 | 7 | -803.79 | -4.87 |
| 19 | 0 | -803.49 | -4.8 |
| 19 | 1 | -803.49 | -4.8 |
| 19 | 2 | -802.63 | -4.74 |
| 19 | 3 | -802.61 | -4.74 |
| 19 | 4 | -802.58 | -4.74 |
| 19 | 5 | -802.58 | -4.74 |
| 19 | 6 | -801.43 | -4.62 |
| 19 | 7 | -801.43 | -4.62 |
| 20 | 0 | -803.47 | -4.8 |
| 20 | 1 | -803.47 | -4.8 |
| 20 | 2 | -803.47 | -4.8 |
| 20 | 3 | -803.47 | -4.8 |
| 20 | 4 | -803.47 | -4.8 |
| 20 | 5 | -803.46 | -4.8 |
| 20 | 6 | -803.46 | -4.8 |
| 20 | 7 | -803.46 | -4.8 |
| 21 | 0 | -803.46 | -4.93 |
| 21 | 1 | -803.46 | -4.93 |
| 21 | 2 | -803.46 | -4.93 |
| 21 | 3 | -803.45 | -4.93 |
| 21 | 4 | -803.06 | -4.95 |
| 21 | 5 | -803.06 | -4.95 |
| 21 | 6 | -803.06 | -4.95 |
| 21 | 7 | -803.05 | -4.95 |
| 22 | 0 | -803.46 | -5.01 |
| 22 | 1 | -803.46 | -5.01 |
| 22 | 2 | -803.46 | -5.01 |
| 22 | 3 | -803.46 | -5.01 |
| 22 | 4 | -803.46 | -5.01 |
| 22 | 5 | -803.46 | -5.01 |
| 22 | 6 | -803.46 | -5.01 |
| 22 | 7 | -803.46 | -5.01 |
| 23 | 0 | -803.4 | -4.8 |
| 23 | 1 | -803.4 | -4.8 |
| 23 | 2 | -803.4 | -4.8 |
| 23 | 3 | -803.4 | -4.8 |
| 23 | 4 | -803.4 | -4.8 |
| 23 | 5 | -803.4 | -4.8 |
| 23 | 6 | -803.4 | -4.8 |
| 23 | 7 | -803.4 | -4.8 |
| 24 | 0 | -803.36 | -4.84 |
| 24 | 1 | -803.36 | -4.84 |
| 24 | 2 | -799.49 | -4.52 |
| 24 | 3 | -799.49 | -4.52 |
| 24 | 4 | -799.49 | -4.52 |
| 24 | 5 | -799.33 | -4.55 |
| 24 | 6 | -792.54 | -4.03 |
| 24 | 7 | -791.99 | -3.91 |
| 25 | 0 | -803.29 | -4.93 |
| 25 | 1 | -803.15 | -4.92 |
| 25 | 2 | -803.14 | -4.92 |
| 26 | 0 | -803.27 | -4.91 |
| 26 | 1 | -803.27 | -4.91 |
| 26 | 2 | -803.27 | -4.91 |
| 26 | 3 | -803.27 | -4.91 |
| 26 | 4 | -803.27 | -4.91 |
| 26 | 5 | -803.27 | -4.91 |
| 26 | 6 | -803.25 | -4.91 |
| 26 | 7 | -803.25 | -4.91 |
| 27 | 0 | -803.12 | -4.88 |
| 27 | 1 | -803.12 | -4.88 |
| 27 | 2 | -802.67 | -4.73 |
| 27 | 3 | -802.67 | -4.73 |
| 27 | 4 | -802.67 | -4.73 |
| 27 | 5 | -802.54 | -4.71 |
| 27 | 6 | -802.54 | -4.71 |
| 27 | 7 | -802.54 | -4.71 |
| 28 | 0 | -803.05 | -4.85 |
| 28 | 1 | -803.05 | -4.85 |
| 28 | 2 | -802.69 | -4.87 |
| 28 | 3 | -802.69 | -4.87 |
| 28 | 4 | -802.69 | -4.87 |
| 28 | 5 | -802.07 | -4.82 |
| 28 | 6 | -802.07 | -4.82 |
| 28 | 7 | -801.68 | -4.84 |
| 29 | 0 | -802.94 | -4.74 |
| 29 | 1 | -802.94 | -4.74 |
| 29 | 2 | -802.94 | -4.74 |
| 29 | 3 | -802.94 | -4.74 |
| 29 | 4 | -802.94 | -4.74 |
| 29 | 5 | -802.94 | -4.74 |
| 29 | 6 | -802.93 | -4.74 |
| 29 | 7 | -802.93 | -4.74 |
| 30 | 0 | -801.91 | -4.73 |
| 31 | 0 | -801.9 | -4.71 |
| 32 | 0 | -801.25 | -4.73 |
| 32 | 0 | -801.25 | -4.73 |

